# Does the Zebra Finch Mating Song Circuit Use Spike Times Efficiently?

**DOI:** 10.1101/2021.03.18.436095

**Authors:** Wilten Nicola, Thomas Robert Newton, Claudia Clopath

## Abstract

Precise and reliable spike times are thought to subserve multiple possible functions, including improving the accuracy of encoding stimuli or behaviours relative to other coding schemes. Indeed, repeating sequences of spikes with sub-millisecond precision exist in nature, such as the synfire chain of spikes in area HVC of the zebra-finch mating-song circuit. Here, we analyzed what impact precise and reliable spikes have on the encoding accuracy for both the zebra-finch and more generic neural circuits using computational modelling. Our results show that neural circuits can use precisely timed spikes to encode signals with a higher-order accuracy than a conventional rate code. Circuits with precisely timed and reliably emitted spikes increase their encoding accuracy linearly with network size, which is the hallmark signature of an efficient code. This qualitatively differs from circuits that employ a rate code which increase their encoding accuracy with the square-root of network size. However, this improved scaling is dependent on the spikes becoming more accurate and more reliable with larger networks. Finally, we discuss how to test this scaling relationship in the zebra mating song circuit using both neural data and song-spectrogram-based recordings while taking advantage of the natural fluctuation in HVC network size due to neurogenesis. The zebra-finch mating-song circuit may represent the most likely candidate system for the use of spike-timing-based, efficient coding strategies in nature.

## Introduction

Our movements, behaviours, and perceptions can display a remarkable consistency from moment to moment, such as when an expert pianist flawlessly performs a recital from memory. This consistency, however, is not a unique characteristic of humans but is a general property of the animal kingdom. One striking example is the imminently reproducible and well-studied mating song of the zebra finch (*Taeniophygia guttata*).

The mating song, which the male zebra finch recites to court females, is learned by a young finch after observing older tutors [Price, 1979, Nottebohm, 1972]. Typically, only a short exposure to the tutor song is required for the song to be internalized. Once internalized and after an extended period of practice, the behaviour becomes learned and can be generated by the student throughout the duration of its life.

This striking reproducibility of a learned behaviour was quickly capitalized on by neuroscientists in an effort to investigate the consistency of learned behaviours. Indeed, we now know that the zebra-finch singing is critically dependent on three primary nuclei post-learning; the HVC (proper name), the Robust Nucleus of the Archopallium (RA), and the hypoglossal nucleus (Figure 1) [Nottebohm et al., 1976, Mooney, 2000, Long et al., 2010, Leonardo and Fee, 2005, Kozhevnikov and Fee, 2007, Hahnloser et al., 2002, Roberts et al., 2012]. During the approximately half-second bout of singing, the HVC neurons which project to area RA (HVC_RA_) fire a precise, chain of spikes where each HVC_RA_ neuron fires a burst at a well-defined moment in time [Hahnloser et al., 2002]. This chain of spikes evenly covers the time-interval of singing, even during the silent intervals between the different segments (or syllables) of a single song [Picardo et al., 2016]. This chain of spikes is also highly reproducible across singing bouts, with individual spikes displaying sub-millisecond precision (Figure 1B, [Leonardo and Fee, 2005, Hahnloser et al., 2002]). At RA, the individual neurons respond to the spectral features of the song and also display highly reproducible spike sequences with sub-millisecond precision [Leonardo and Fee, 2005]. This is one of the final command signals to produce singing through the vocal organ with the hypoglossal nucleus acting as the final relay. Thus, the core neural architecture at the heart of this highly reproducible and precise behaviour is a highly reproducible and precise sequence of spikes.

**Figure 1:**
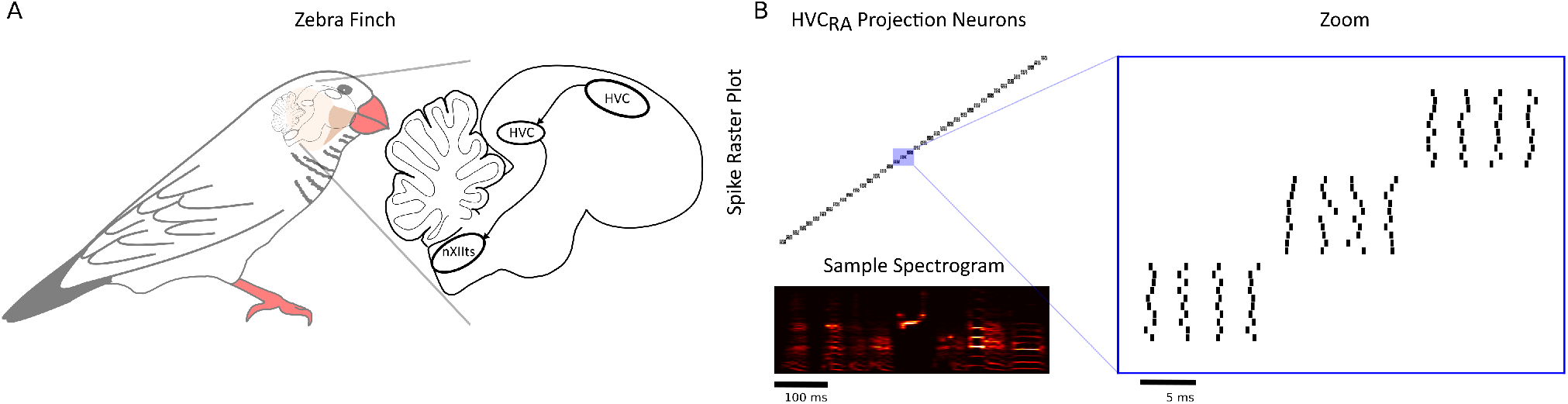
**(A)** The Zebra-Finch mating song circuit consists of areas HVC, RA, and the hypoglossal nucleus (nxIIts). These are the minimum areas required to produce the mating song in an adult bird. **(B)** Area HVC contains neurons which project to RA (HVC_RA_). These neurons fire a classical syn-fire chain of spikes which uniformly cover time during a song-replay. The spikes in HVC repeat during each song replay with sub-millisecond precision [Hahnloser et al., 2002, Leonardo and Fee, 2005].

However, precisely timed spike sequences as observed in HVC_RA_ are perhaps the exception, rather than the rule, in the animal kingdom, despite the theoretical advantages that spike times may have [Bohte, 2004, VanRullen et al., 2005, Panzeri et al., 2001, Gutkin et al., 2003, Borst and Theunissen, 1999, Gütig, 2014, Tully et al., 2016, Quiroga and Panzeri, 2009, Victor and Purpura, 1996] Indeed, in vivo recordings from neurons in pre-motor areas of other animals do not typically demonstrate the precise timing of spikes as found in HVC [Wang et al., 2018, Chestek et al., 2007]. In fact, rate-codes, where the number of spikes per unit time encodes information are reliably observed. If other pre-motor areas use a rate-code for the reproducibility of behaviours, what advantage does the zebra finch obtain in precisely controlling the timing of spikes?

In this study, we considered the question of what benefit precise-spike times provide over a rate-code from a neural coding perspective. By using synthetically constructed spike-trains and mathematical analysis, we found that reliable and accurate spikes do have a qualitative difference over rate-codes in the accuracy of encoding behaviours and signals: the error in encoding a behaviour is inversely proportional to the network size. This scaling-relationship, where a network can double its size to halve the error, is in strike contrast to a rate-code where a network has to quadruple in size to halve the error, and is therefore a hallmark for an efficient code [Barlow et al., 1961, Denève and Machens, 2016a, Denève and Machens, 2016b, Boerlin et al., 2013, Schwemmer et al., 2015]. We both mathematically derived and tested these findings in numerical simulations with synthetically generated spike-trains and recorded song-bird spectrograms. This linear scaling-relationship also holds when the spike-times are no longer perfectly accurate and reliable, so long as the timing of spikes and their reliability in being emitted by a neuron increases along with the size of a network. Finally, we discuss how to test the scaling-relationship experimentally by either exploiting the natural increase in the HVC network size (via neurogenesis) or through potential optogenetic, or neural-ablation based perturbations.

## Results

Before we consider how precise spike-times impact the neural code, we need to formally define neural encoding and decoding. As an organism performs a behaviour, or perceives a stimulus, which we will define with the function of time *x*(*t*), spikes are emitted as a result of the stimuli or in pursuit of enacting the behaviour. The relationship between the spikes and behaviour can be deduced by constructing a decoder, *ϕ^x^* which transforms the spike train ***r***(*t*) into an approximation of the behaviour or stimulus 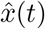:

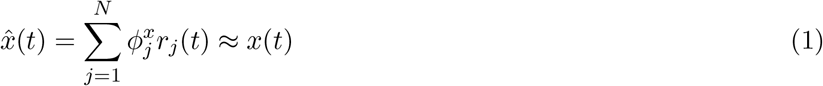

Typically, the decoder is constructed by using one or a subset of repetitions of the behaviour or stimulus (and spikes) and later tested with repetitions that were not used to construct the initial decoder. When 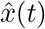 can accurately decode out *x*(*t*), then the spike train ***r***(*t*) = [*r*_1_(*t*), *r*_2_(*t*),… *r_N_*(*t*)] generated for the *N* neurons is encoding some information about the stimulus or behaviour *x*(*t*).

The very existence of neural encoding and decoding, therefore, raises the question of how accurate is the neural code? The Root-Mean-Squared Error (RMSE) is commonly used as the metric to measure this accuracy [Denève and Machens, 2016b, Boerlin et al., 2013, Schwemmer et al., 2015]. For example, if we consider the audio-time series of a zebra-finches singing as *x*(*t*), then the RMSE measures the difference between the neurally decoded song 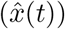 and the recording with an RMSE of 0 indicating perfect decoding accuracy.

The core result of this research is in demonstrating that for precisely timed sequences of spikes, such as those fired by HVC_RA_ neurons or pyramidal neurons in RA in the zebra-finch mating song circuit, the RMSE decreases linearly with the number of neurons:

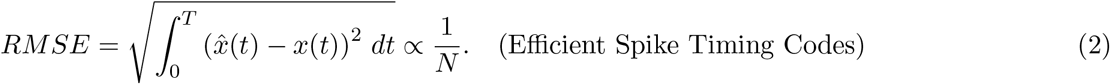

This is qualitatively different from and in strike contrast to rate codes, where the RMSE decreases with the square root of the number of neurons [Denève and Machens, 2016b, Boerlin et al., 2013, Schwemmer et al., 2015]:

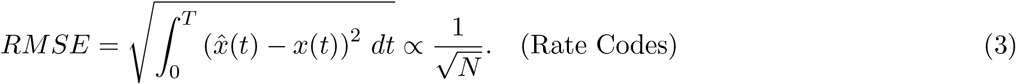

### Timing Codes Scale Linearly with the Network Size in the Root-Mean-Squared-Error (RMSE) under Optimal Linear Decoding

Given the observed spike-timing precision in the mating song circuit, we naturally wondered what the impact of precise spike-timing would have on the RMSE of decoded behaviours or stimuli. We first constructed a simplified model of the mating song circuit with synthetically constructed spike trains that mimic the statistics of HVC_RA_ neurons (Figure 2A-B). In particular, each HVC_RA_ neuron fires a single burst of spikes, and the bursts are uniformly distributed over the entire duration of the song (Figure 2C). A decoder (*ϕ^x^*) is then constructed using a song-recording from a real song (see Materials and Methods, [Nicola and Clopath, 2017]). We found that larger networks (increasing N) of HVC_RA_ neurons resulted in more accurate decoding (Figure 2D). Critically, we found that for sufficiently large networks, further increases lead to a nearly linear decrease in the RMSE with network size (RMSE ∝ *N*^−0.955^). This is in stark contrast to what would be expected with a rate code, where the RSME typically decreases with the square root of the network size (RMSE ∝ *N*^−0.5^).

**Figure 2:**
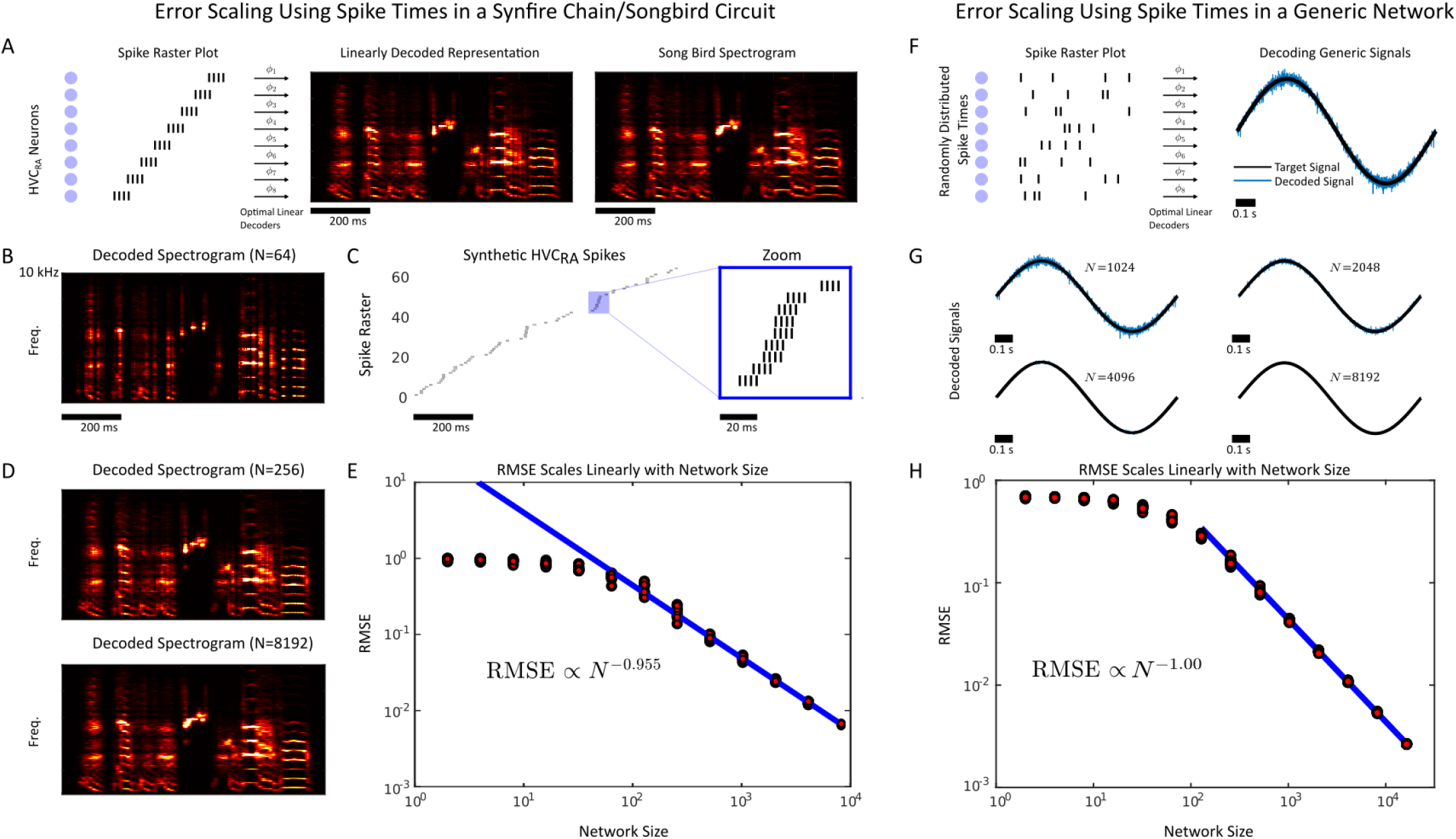
**(A)** Synthetic HVC_RA_ synfire chain spike sequences are created with perfectly precise and reliable spikes. These spikes are used to decode a sample-spectrogram from a zebra-finch mating-song recording with an optimal linear decoder. **(B)** A decoded spectrogram using *N* = 64 HVC_RA_ neurons. **(C)** The HVC_RA_ spikes consist of stereotypes bursts of 4 spikes, shifted randomly in time to cover the duration of song replay **(D)** Larger networks encode the song-spectrogram with increased accuracy. **(E)** The accuracy, as measured by the Root-Mean-Squared Error (RMSE) scales linearly with the network size. **(F)** A generic, randomly generated spike sequence with perfectly precise and reliable spike times are used to linearly decode a generic signal (sinusoidal oscillator). **(G)** Larger networks encode the sinusoidal signal with increasing accuracy **(H)** RMSE decreases linearly with the network size, as in the HVC_RA_ network demonstrating that *N*^−1^ scaling in the RMSE is a generic phenomenon which is a result of having precise and reliable spikes.

Next, we sought to determine how general this result was. Was the linear decrease with network size somehow brought on by isolated bursts representing a synfire chain [Herrmann et al., 1995, Abeles, 2012], or was this a more general phenomenon due to the precision in the spike times? To investigate this further, we constructed synthetic spike trains where each neuron could fire a variable amount of spikes (drawn from a Poisson distribution) that were reliable and precisely timed from trial to trial (Figure 2F). We again constructed decoders that could decode out a target signal, in this case, a simple oscillator (Figure 2F). We found that the RMSE also decreased linearly (RMSE ∝ *N*^−1.00^) with the network size for sufficiently large networks, indicating that its spike-timing precision that results in *N*^−1^ RMSE error scaling, rather than the synfire-chain nature of the spikes in the HVC_RA_ circuit.

Collectively, these numerical results seem to imply that precise-spike times reliably lead to *N*^−1^ scaling in the RMSE. Thus, we investigated under what conditions this result was guaranteed to hold (Supplementary Appendix, Supplementary Figure 1). We found that the linear decrease in the RMSE with network size is a general, mathematical result. So long as the spikes emitted by neurons are precise from trial-to-trial, and sufficiently dense in time, the vast majority of emitted spike trains lead to *RMSE* ∝ *N*^−1^ for sufficiently large network sizes.

### The Precision and Reliability of Spike Times Must Increase with the Network Size to Preserve Linear Error Scaling

While spike-times in HVC_RA_ are precise, they are not perfectly precise. From trial-to-trial, the spike times often display sub-millisecond levels of jitter [Hahnloser et al., 2002, Leonardo and Fee, 2005]. Further, neurons can also fail in firing spikes owing to unreliable synapses. Thus, we next considered just how precise spikes have to be to lead to *N*^−1^ scaling in the RSME.

First, we considered spike-jitter from trial-to-trial by constructing a decoder with initially precise spikes. Then, we perturbed the spike times randomly and reapplied the decoder on subsequent trials, and measured the RMSE (Figure 3A). We found that if the standard deviation of the jitter (*σ*) was fixed, the *N*^−1^ error scaling in the RMSE was no longer present (RMSE ∝ *N*^−0.187^).

**Figure 3:**
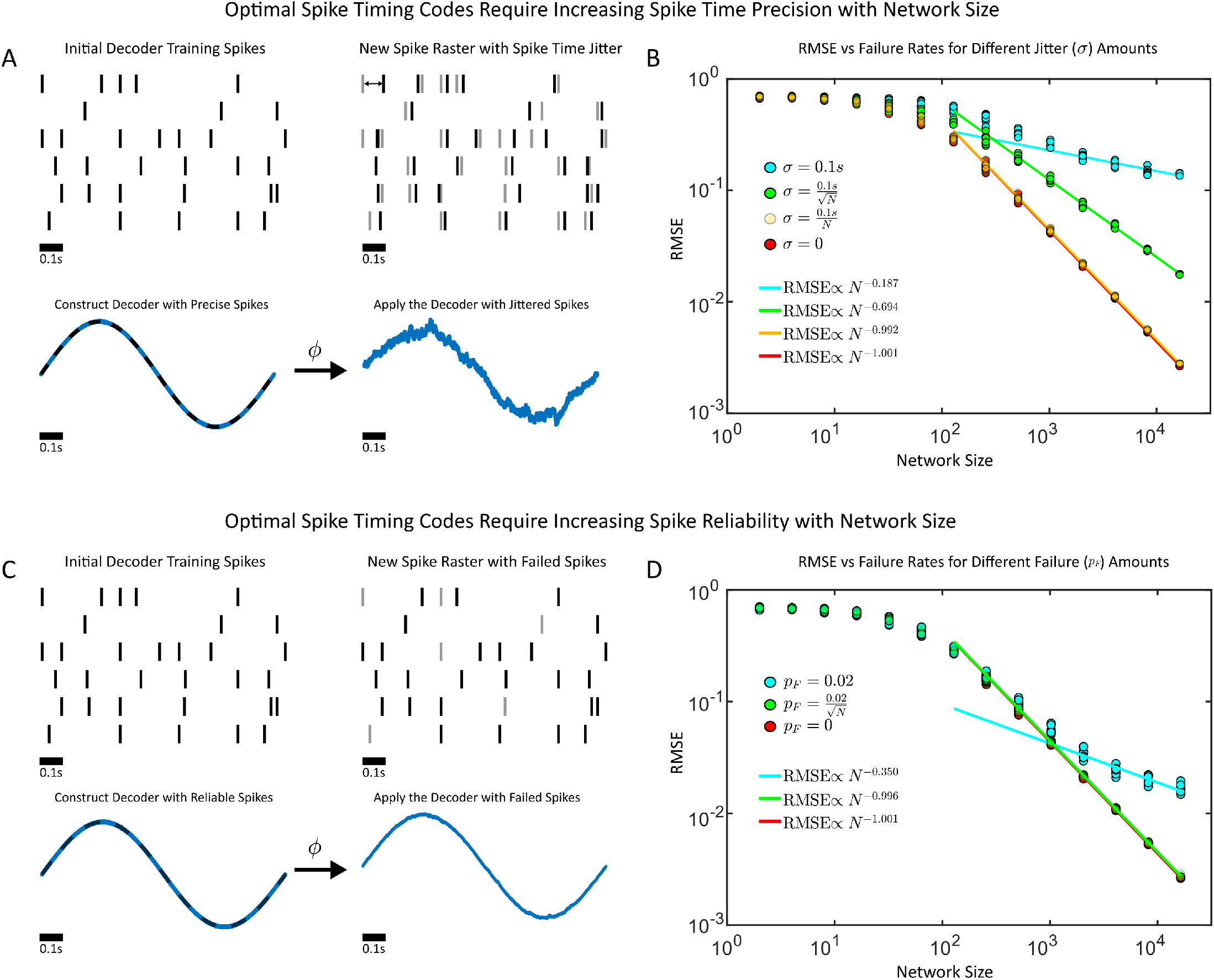
**(A)** Synthetic spike trains are generated and used to train an optimal decoder on a simple sinusoidal oscillator. The spike trains are then jittered with and the decoder is reapplied, with the resulting root mean squared error (RMSE) measured. **(B)** The RMSE as a function of the network sized (*N*) for differing jitter amounts *σ*. **(C)** Synthetic spike trains are generated and decoders are trained as in (A), only now a random subset of spikes fail with *p_F_* denoting the spike failure probability. A *p_F_* = 1 implies that all spikes fail, while a *p_F_* = 0 implies that the spikes are perfectly reliable. **(D)** The RMSE as a function of the network sized (*N*) for differing spike failure amounts *p_F_*.

If a fixed amount of imprecision would destroy the *N*^−1^ scaling, could networks increase their spike timing precision with network size to restore it? To investigate this, we considered the possibility that spikes might become more precisely timed with increasing network sizes by having the jitter decrease with larger networks (either 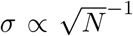 or *σ* ∝ *N*^−1^, Figure 3B). We found that if the spike-times became more precise linearly (*σ* ∝ *N*^−1^) with the network size, this was sufficient to restore the *N*^−1^ error scaling in the RSME (RMSE ∝ *N*^−0.992^).

Finally, we considered the case where spikes might abruptly fail (Figure 3C). We again found that if the probability of spike failure (*p_F_*) was fixed and independent of the network size, the *N*^−1^ scaling in the RMSE would no longer be present (RMSE ∝ *N*^−0.350^). However, once again, if the spikes are less likely to fail with larger networks 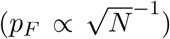, the *N*^−1^ scaling in the RMSE would be restored (RMSE ∝ *N*^−0.996^). These results were qualitatively and quantitatively similar to the case where spikes were randomly emitted (rather than randomly failed, Supplementary Figure 2) Thus, spike-timing imprecision or spike-unreliability are themselves not sufficient to eliminate the RMSE ∝ *N*^−1^ scaling, but the spikes must become more reliable and more precise with progressively larger network sizes to maintain *N*^−1^ RMSE scaling.

### The songbird circuit can maintain 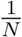 RMSE scaling with if HVC_RA_ spikes become more precise with increasing network size

If spike-timing imprecision could eliminate the *RMSE* ∝ *N*^−1^ relationship, could the highly precise HVC_RA_ spikes still display an *N*^−1^ RMSE scaling? To test this hypothesis, we constructed increasingly larger networks of synthetic spike trains and decoded the spectrogram of a sample song with imprecise spikes (Figure 4A). The spikes used for the decoding were jittered versions (*σ* = 0.3 ms) of the spikes used to construct the initial decoder. We found that for large networks, 2^10^ < *N* < 2^15^, the linear scaling in the RMSE was no longer present with the resulting slope in the log-log plot of the RMSE versus network size having a numerical value of −0.43, which is closer to (and worse than) the slope expected from a rate code (slope = −0.5, Figure 4B-C)

**Figure 4:**
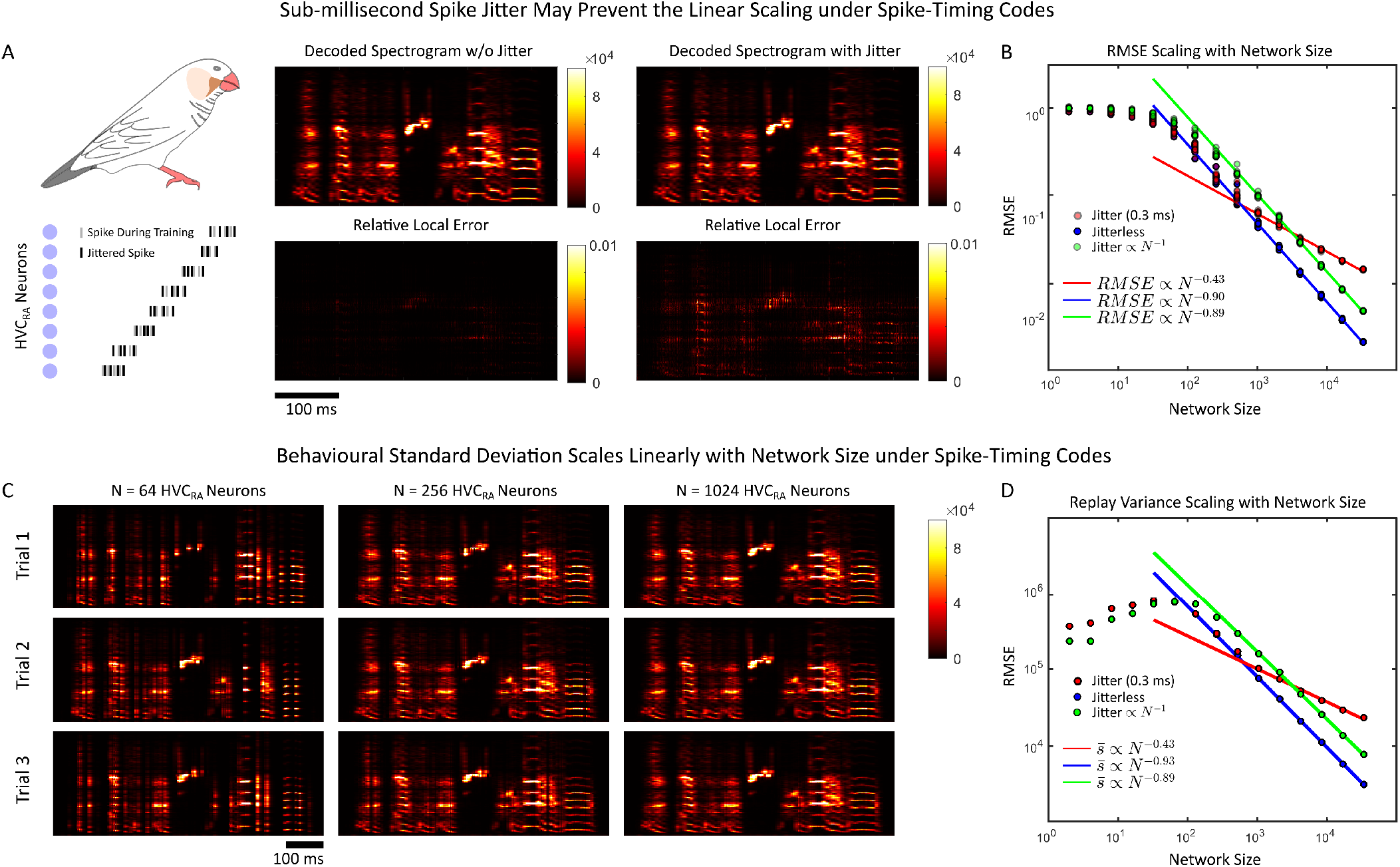
**(A)** Synthetic HVC_RA_ synfire chain spike sequences are created and used to train a decoder to decode out a sample spectrogram. These spikes are then jittered with varying amounts of jitter with the decoder reapplied and the RMSE measured. **(B)** Root Mean Squared Error (RMSE) between the song spectrogram and the decoded output as a function of the network size (*N*) and with varying amounts of jitter. Sub-millisecond jitter can destroy the ≈ *N*^−1^ scaling in the RMSE while increasingly precise spike-times (jitter ∝ *N*^−1^) can restore it. **(C)** Sample decoded spectrograms for different values of the network size for the fixed jitter *σ* = 0.3 ms case. **(D)** The spectrogram standard deviation 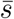 (see Materials and Methods) displays identical scaling relationships as the RMSE.

Our previous simulations (Figure 3) implied that the RMSE ∝ *N*^−1^ scaling relationship could be rescued under one condition: spikes becoming more precise with larger networks. To test if this principle still holds in the songbird circuit, we linearly decreased the spike jitter standard deviation *σ* with larger networks *σ* ∝ *N*^−1^, so that a doubling of the network size also doubled the precision of spike times. We found that this reliably restored the nearly linear scaling with network size of the RMSE (*N*^−0.89^).

While the RMSE is important for computational studies, it is difficult for an experimentalist to actually utilize and measure for a simple reason: they do not have access to the actual signal a circuit intends to represent, only the output of the circuit itself. Indeed, the RMSE represents the accuracy of a behaviour once that behaviour is known. Thus, we tested whether the precision of a behaviour also increased with network size by measuring the standard deviation of the decoded spectrograms (Figure 4D). Fortunately, we found that the replay variability (see Materials and Methods for definition) of the spectrograms displayed very similar scaling relationships (slopes of −0.43, −0.93, and −0.89) as the RMSE (slopes of −0.43, −0.9, and −0.89) for all conditions (*σ* = 0.3 ms, *σ* = 0, *σ* ∝ *N*^−1^). This result allows an experimentalist to record a small number of repetitions (10 in this case) of animal singing, and use the stereotypy of a song as a proxy for the RMSE in measuring these scaling relationships.

## Discussion

Precisely timed spikes have long been hypothesized to somehow encode information [Bohte, 2004, VanRullen et al., 2005, Panzeri et al., 2001, Gutkin et al., 2003, Borst and Theunissen, 1999, Gütig, 2014, Tully et al., 2016, Quiroga and Panzeri, 2009, Victor and Purpura, 1996]. Here, we explored the hypothesis that spike timing precision and spike reliability might be specifically used to improve the accuracy of encoding in a way that is qualitatively different from a rate code. The error in decoding any behaviour or sensory stimulus decreases linearly (rather than square-root) with the network size. We found that with synthetically generated spike trains that represent the precisely timed HVC_RA_ neurons in the zebra-finch circuit, or in more generic precisely timed circuits display a linear decrease in the root-mean-squared-error (RMSE) with the network size. If the variance in the spike times remained fixed, this linear relationship would be reduced to either rate-code (RMSE ∝ *N*^−0.5^) or even sub-rate code error scaling levels. If, however, spike reliability and spike timing precision increased with the network size, then the inverse linear relationship between RMSE and network size would be retained. Finally, the RMSE can be entirely replaced as a proxy with the standard deviation or stereotypy of an empirically measured behaviour, allowing experimentalists to easily test the impacts of spike-timing precision on the precision of behaviours without knowing the true intended ‘‘target” behaviour.

The zebra finch mating song circuit is uniquely positioned to test long-standing hypotheses about how the precision in reproducing behaviours is determined by the number of neurons controlling said behaviour. First, the circuit controls a stereotyped behaviour that is readily elicited and easily recorded with a microphone in head-fixed animals. Second, new HVC_RA_ neurons naturally form through the process of neurogenesis [Walton et al., 2012, Kirn and Nottebohm, 1993, Pytte et al., 2012, Brenowitz and Larson, 2015, Pytte et al., 2007, Nordeen and Nordeen, 1988] in HVC, and roughly double in number from an average of around 40, 000 HVC_RA_ neurons in the first year of life, to around 80,000 HVC_RA_ neurons by year 11 (see Figure 2 in [Walton et al., 2012], Figure 5). The core prediction of our work is that under efficient spike-timing codes, singing bouts will increase in their precision, or equivalently, decrease in their standard deviation linearly with the size of the network (Figure 5). The precise trend comparing the behavioural precision with the network size of HVC_RA_ neurons has not been currently ascertained. Tantalizingly, there are a pair of studies that separately imply that the total number of HVC_RA_ neurons increases with age ([Walton et al., 2012]) and that the singing precision also increases with age ([Pytte et al., 2007]). However, the authors in [Pytte et al., 2007] postulated that the increase in behavioural precision was inversely related to the rate of neurogenesis, rather than the overall number of HVC_RA_ neurons.

**Figure 5:**
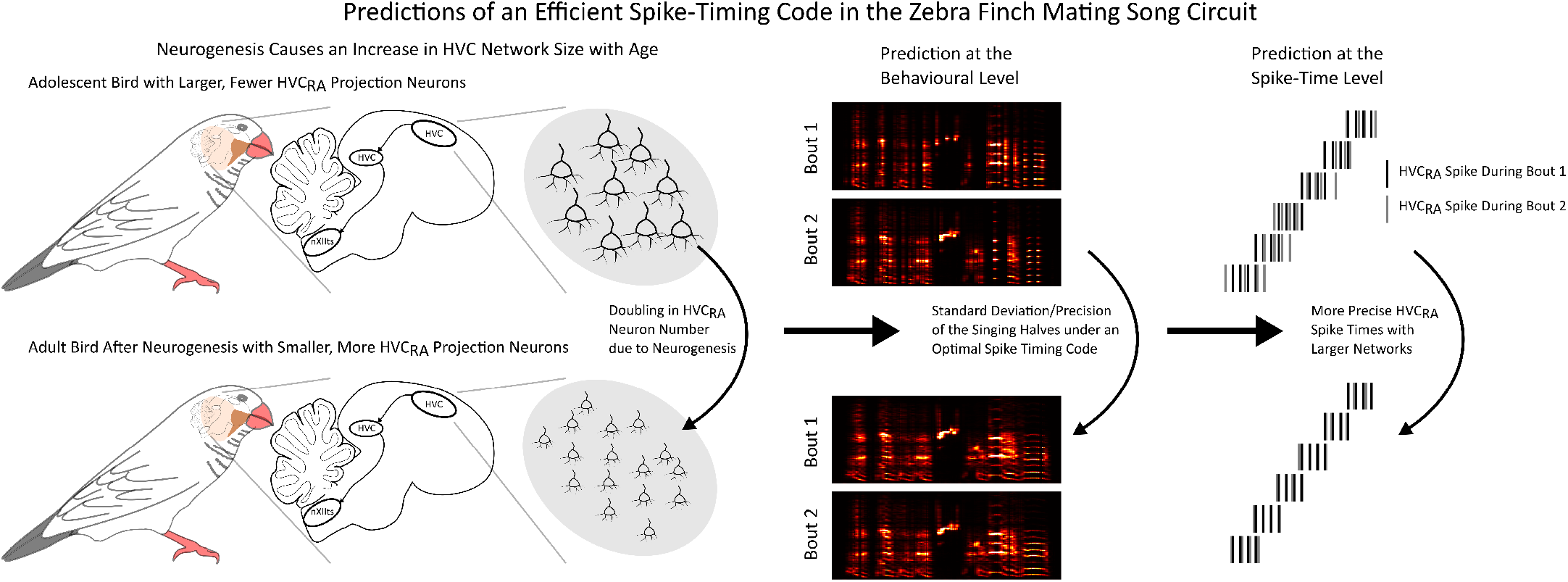
The core prediction of an effcient spike timing code in the zebra Finch mating song circuit. An increase in network size (in this case a doubling) due to neurogenesis leads to a proportional increase in behavioural stereotypy/precision (or decrease in the behavioural standard deviation), and an increase in the precision of spike times in HVC_RA_ neurons

As an alternative to naturally allowing the HVC_RA_ neurons to increase in number with neurogenesis, one can selectively inhibit HVC_RA_ neurons with optogenetics or ablate a fixed proportion of the HVC nucleus entirely [Roberts et al., 2012]. This would deactivate a random subset of HVC_RA_ neurons. After the animal is given a suitable amount of time to recover and relearn from the deactivation of said neurons, the resulting circuit should produce the identical song, but with fewer HVC_RA_ neurons. This recovery period should also be smaller than the (very slow) time-scale of HVC_RA_ neurogenesis. Here, we would expect the behavioural precision to decrease linearly with the proportion of neurons deactivated.

Further, we remark that precise and reliably emitted spikes are sufficient to generate a linear decrease of the RMSE with network size, however, this is not the only possibility. In fact, linear error scaling was previously predicted under efficient spike-time coding schemes [Denève and Machens, 2016b, Boerlin et al., 2013, Schwemmer et al., 2015, Denève and Machens, 2016a, Brendel et al., 2020]. Here, neurons explicitly code the error in the representation of a stimulus, behaviour, or internal dynamic state with their voltage. When the error reaches a critical threshold, a neuron fires a spike to explicitly reset the error to 0. The end result is a network that also produces a linear scaling of the RMSE with network size. However, owing to the explicit error correction mechanism in these circuits, the spike times for individual neurons are not precisely timed across multiple trials, but are precisely timed to reset the error in a representation [Denève and Machens, 2016b, Boerlin et al., 2013, Schwemmer et al., 2015, Denève and Machens, 2016a, Brendel et al., 2020]. Here, we suggest an alternate mechanism for *N*^−1^ error scaling in a circuit which, at face values, displays the expected properties of an efficient spike-timing code: behavioural stereotypy with precisely timed spikes.

Finally, efficient coding in the mating song circuit has been considered in one other modelling study where rate-based artificial neural networks were trained to reproduce song-bird spectrograms [Blättler and Hahnloser, 2011]. Our work differs here in two important ways. First, we analyze the specific contribution of precise spike times to the production of song. Second, we translate these findings into an efficient coding hypothesis for the singing behaviours precision and accuracy [Denève and Machens, 2016b, Boerlin et al., 2013, Schwemmer et al., 2015, Denève and Machens, 2016a, Brendel et al., 2020].

This theoretical work demonstrates that it is possible for a neural circuit to efficiently code a behaviour (or a stimulus) with precisely timed and reliably emitted spikes. Under efficient spike-timing codes, a network can double its size to double the precision and accuracy of encoding. The zebra-finch mating song circuit presents what is arguably the best opportunity to test for the existence of an efficient code in nature, or if we are likely always limited in encoding accuracy due to the imprecise and unreliable nature of neuronal spiking.

## Materials and Methods

### Generating Synthetic Spike Trains

Each network we consider here constitutes *N* synthetically generated spike trains, with each spike train representing a single neuron. This allows us to create synthetic spike times, *t_ji_* which corresponds to the *i*th spike fired by the *j*th neuron that could have their spike-timing precision and spike-reliability controlled. These spikes were then filtered with a synaptic filter:

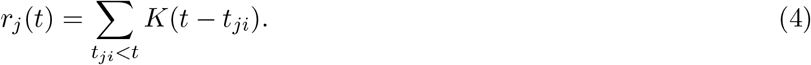

The filtering function, *K*(*t*) was taken to be a single-exponential filter for all numerical simulations:

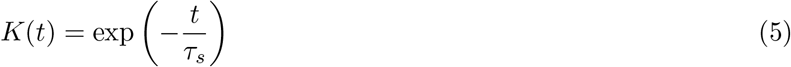

where *τ_s_* = 5 ms was used as the synaptic filter, approximating the time constant of AMPA synapses. The generation of spike times *t_ji_* is described in greater detail below.

### HVC_RA_ Spike Train

Each HVC_RA_ neuron consisted of a synthetically generated spike train consisting of 4 spikes in an isolated burst. The spikes were separated by 3 ms inter-spike-intervals, with the initial spike-time being uniformly distributed on the interval [0,*T*], where *T* = 0.88 seconds is the duration of the song-recording. To vary the network size *N*, the networks were successively doubled in size from *N* = 2^1^ to *N* = 2^15^ = 32768. The networks in Figure 2 and Figure 4 were simulated 10 times for a fixed *N*, with different seeds in each network, thereby generating different starting times for each HVC_RA_ burst.

To jitter the spikes in Figure 4, each spike time *t_ji_* was randomly perturbed by *ϵ_ji_*, where *ϵ_ji_* was a normally distributed random variable with mean 0, and standard deviation, *σ*. The values of *σ* vary within figures with perfectly precise spikes (*σ* = 0 milliseconds), spikes with sub-millisecond precision (*σ* = 0.3 milliseconds), and spikes that become increasingly precise with larger network sizes 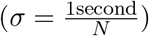.

### Poisson Generated Spike Trains

The randomly generated spikes in Figure 2F-H and 3 were generated from a Poisson process with firing rate *ν* = 2 Hz, for successively larger networks which were doubled in size from *N* = 2^1^ to *N* = 2^14^. As in the songbird example, the spikes were perturbed in time with a normally distributed random variable with mean 0 and standard deviation *σ* (see Figure 3B for *σ* values). To implement spike-failure, decoders were first constructed with all spikes generated. These decoders remained fixed, even after the spike times or spike reliability is altered.

Then, the fixed decoders were applied to the same spike trains but with each spike having a probability of *p_F_* of failure (see Figure 4D for *p_F_* values). To implement spike interference, spikes were added randomly to neurons in time with probability *p_I_*. The total number of spikes added was *p_I_n_spikes_* where *n_spikes_* was the number of spikes generated in the nominal, reliable spiking case. The added spikes were distributed uniformly in time, and across neurons.

### Constructing Linear Decoders

Linear decoders were constructed by first generating filtered spike-trains that contained no jitter or failure. Then the solution to the optimal linear decoder for a given signal *x*(*t*) is:

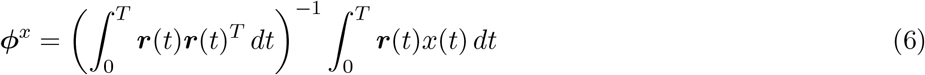

The decoder can then be applied to new spike trains to decode 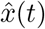 when these new spikes display either imprecision in their spike times or the failure of spike emission relative to the initial spike train used to construct *ϕ^x^*. For the simple sinusoidal example, *x*(*t*) = sin(2*πt*), with the approximation 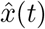 given by:

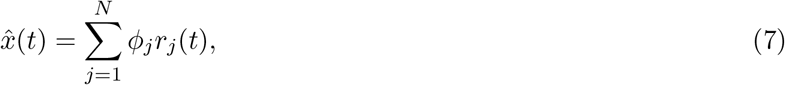

while for the spectrogram, each frequency component of the song has its own decoder:

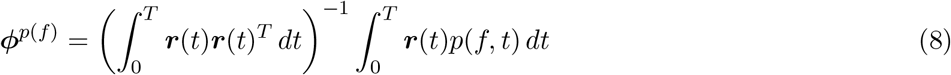

The term *p*(*f, t*) is the power of the frequency component *f* at time *t* in the spectrogram, as defined in [Nicola and Clopath, 2017], while the decoder component *ϕ*^*p*(*f*)^ is the optimal decoder for the power frequency component *p*(*f, t*), yielding the approximation:

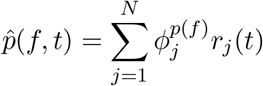

The frequency range considered varies from a low of *f* = 172.27*Hz* to a high of *f* = 10 kHz, discretized with 229 points uniformly distributed points, thereby making *ϕ*^*p*(*f*)^ a *N* × 229 dimensional decoder.

### Measuring the Root Mean-Squared Error

The RMSE for the decoded oscillators in Figures 2F-H and 3 is given by:

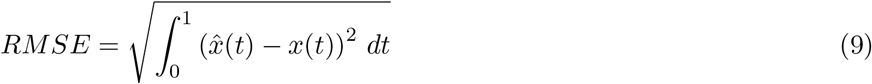

where *x*(*t*) = sin(2π*t*), and 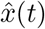 is the decoded approximation to *x*(*t*).

The RMSE for the decoded spectrogram is computed with:

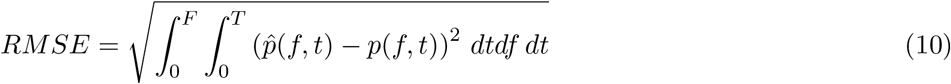

where *p*(*f, t*) is the value of the song spectrogram at frequency *f* and time *t* over the range *f* = 0 to *F* = 10 kHz and from time *t* = 0 time *T* = 0.88 s (song duration). For each value of *N*, 10 trials (corresponding to different decoders) were used to obtain the average value of the RMSE for that fixed value of *N*. The linear fits to the RMSE were constructed on the log-log scale with the MATLAB polyfit function, for sufficiently large networks (*N* ≥ 2^10^).

### Measuring the Stereotypy in the Zebra Finch Networks

To estimate the variability from song-replay to song-replay, we computed the integrated standard deviation, 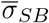:

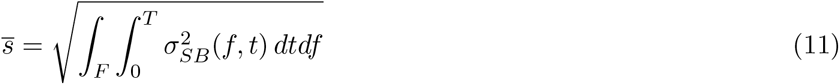

where

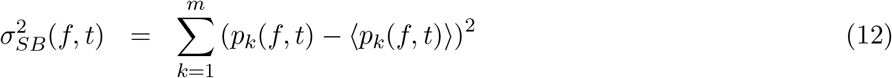

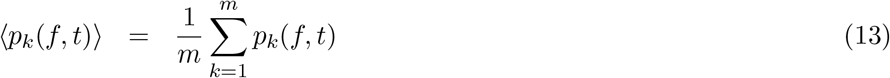

where 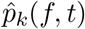 denotes the *k*th trials decoded spectrogram for *k* = 1, 2,… 10.

## Supplementary Materials

### Appendix: Error Scaling with Precise Spike Times

Here, we will show for a general class of spike trains with precisely timed spikes generated by *N* neurons, the following result holds:

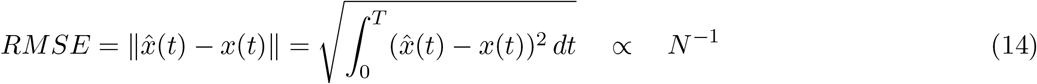

where 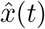 is the neurally decoded approximation to *x*(*t*), given by the following:

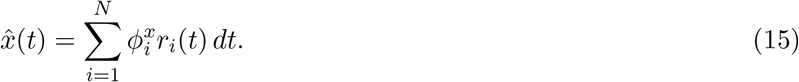

where 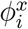 is the optimal decoder for *x*(*t*).

The term *r_i_*(*t*) is the filtered sequence of spike times for neuron *i*:

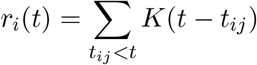

where *K*(*t*) is some filtering function.

The derivation of (14) will be broken into two steps. In the first step, we will prove (14) for the case where each neuron fires a single spike, and the spikes are uniformly spread over the interval [0, *T*]. In the second step, we will prove that the same result holds for more general spike rasters by using linear transformations to determine when a general spike raster can be transformed into the evenly distributed one. The following derivation follows largely from classical approaches from functional analysis, the theory of function approximation, and the Simple Function Approximation Theorem [Davidson and Donsig, 2002, Rudin, 1976]. The 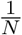 in the RMSE scaling is indeed the “unavoidable discretization error” stated by [Denève and Machens, 2016a], and numerically demonstrated in [Denève and Machens, 2016a] (Figure 1a, the regular-rate code).

### Step 1: Evenly Distributed Spikes

Suppose the spike train fired by the *N* neurons is evenly distributed over an interval [0,*T*], reminiscent of the HVC_RA_ projection neurons [Hahnloser et al., 2002]. Thus, *N* neurons fire at successive 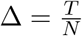 where *T* is the total duration of the signal to be approximated, *x*(*t*). Each neuron fires at times *t_j_* = *t*_*j*–1_ + Δ Suppose further more that we decode each spike with a box filter:

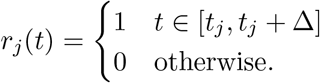

In order to approximate the signal *x*(*t*), on an interval [0, *T*], we need to determine what the decoders 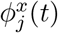 for each neuron *j*. The decoders are easily resolved as the intervals [*t*_*j*–1_, *t_j_*] are non-overlapping, and with box-filtering, the decoded spikes *r_j_*(*t*) are orthogonal. This immediately yields the following decomposition of the approximant:

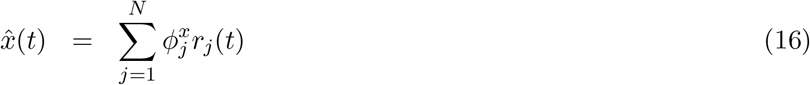

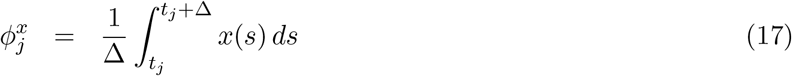

The formula for the decoders is easily derived when one considers the orthogonality of the spike train. Note that if we determine the order of error in Δ, then with Δ = *T/N* we determine how the error scales with the network size. The squared error is thus:

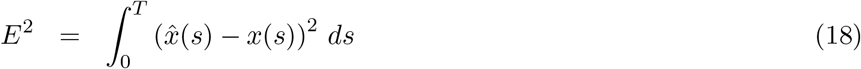

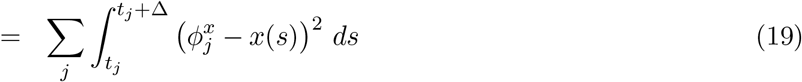

Now, if we can analytically determine or bound the integral

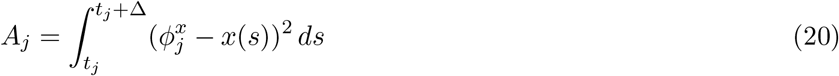

then we can bound the error in equation (18). First, note that by the mean-value theorem for integrals, on the interval [*t_j_, t_j_* + Δ] there exists a *c_j_* ∈ [*t_j_, t_j_* + Δ]

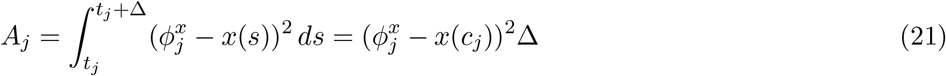

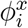 is the mean value *x*(*t*) over [*t_j_, t_j_* + Δ], by definition, thus by the intermediate value theorem, there must exist some 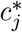 such that 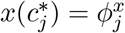, then we have

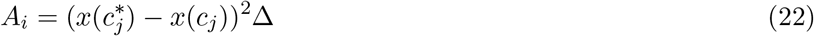

We use the mean-value theorem for derivatives on the smaller interval 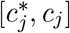. We can assume without loss of generality that 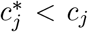, as the opposite case is entirely identical. The mean-value theorem tells us there exists some 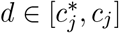 such that:

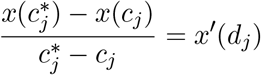

and thus:

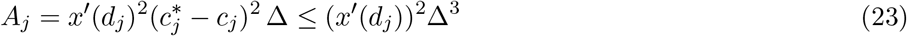

where the inequality comes from the fact that the interval 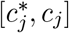 lies within [*t_j_, t_j_* + Δ] and thus 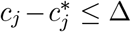. This yields the following:

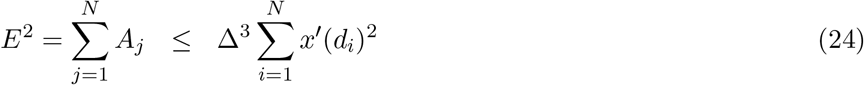

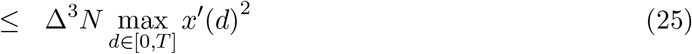

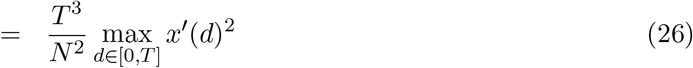

Thus, we have the following:

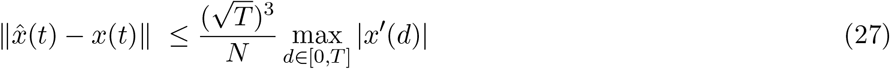

Result (27) implies that for uniformly distributed, precisely timed spikes, the RMSE in approximating a function is inversely proportional to the network size. Doubling the network size halves error, unlike in a conventional rate-code, where the network must quadruple in size to halve the RMSE.

### Step 2: Use Step 1 To Derive Result for More General Spike Trains

More generally, we will assume that the spikes are not uniformly distributed, however the time intervals defined above, [*t_j_, t_j_* + Δ], *j* = 1, 2,… *N* remain, and we still consider a network of *N* neurons. Thus, the following matrix emerges:

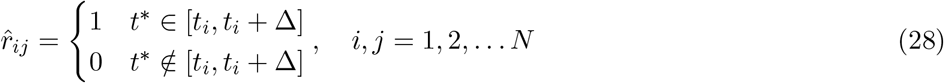

where element (*i, j*) of 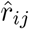 is 1 if neuron *j* fires a spike (*t*\*) in the ith time interval, and 0 otherwise. Finally, we will assume that 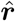 is an invertible matrix, or equivalently, the rank of 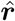 is *N*. Note that if we consider the matrix generated for the uniformly distributed spiking case above (***r***), then ***r*** = ***I**_N_* where ***I**_N_* is the *N* × *N* identity matrix.

Then, if the matrix 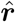 is invertible, consider the decoder defined by:

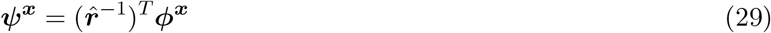

where ***ϕ^x^*** is the same decoder as in the uniformly distributed spiking case considered above. Applying *ψ^x^* to 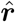 yields

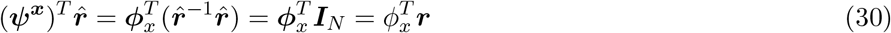

which restores the uniformly distributed spiking approximation in Step 1.

Now, consider the optimal linear decoder for the spike train 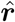, as given by 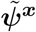. Then, we have the following:

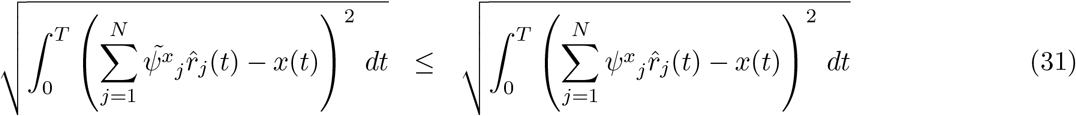

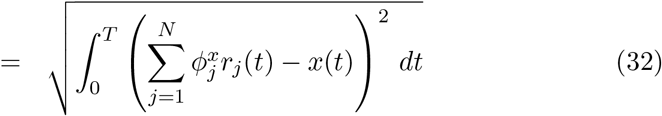

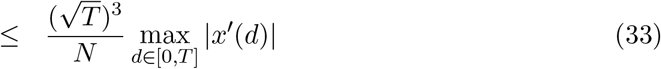

Thus, result (33) shows that any invertible spike-train is bounded by an *O*(*N*^−1^) error. This is in principle most randomly generated spike trains for sufficiently large *N* as these matrices are highly likely to be full-rank (see for example [Tao and Vu, 2007, Bourgain et al., 2010].

As a final comment, we note that this result has an implication for trained spiking neural networks with linear decoder-construction based approaches [Nicola and Clopath, 2017, Nicola and Clopath, 2019, Abbott et al., 2016, DePasquale et al., 2018, Thalmeier et al., 2016, Eliasmith and Anderson, 2003, Gilra and Gerstner, 2017, Zenke and Ganguli, 2018, Chestek et al., 2007, Florian, 2012]. In particular, if the timing of spikes are stabilized to be precisely reproducible between training and testing phases, then *O*(*N*^−1^) convergence is mathematically guaranteed. This, however, is not a necessary criterion, as under error-correcting spike-based codes (such as [Denève and Machens, 2016b, Boerlin et al., 2013, Schwemmer et al., 2015]), *O*(*N*^−1^) convergence can still be achieved without precise spike times.

**Supplementary Figure 1:**
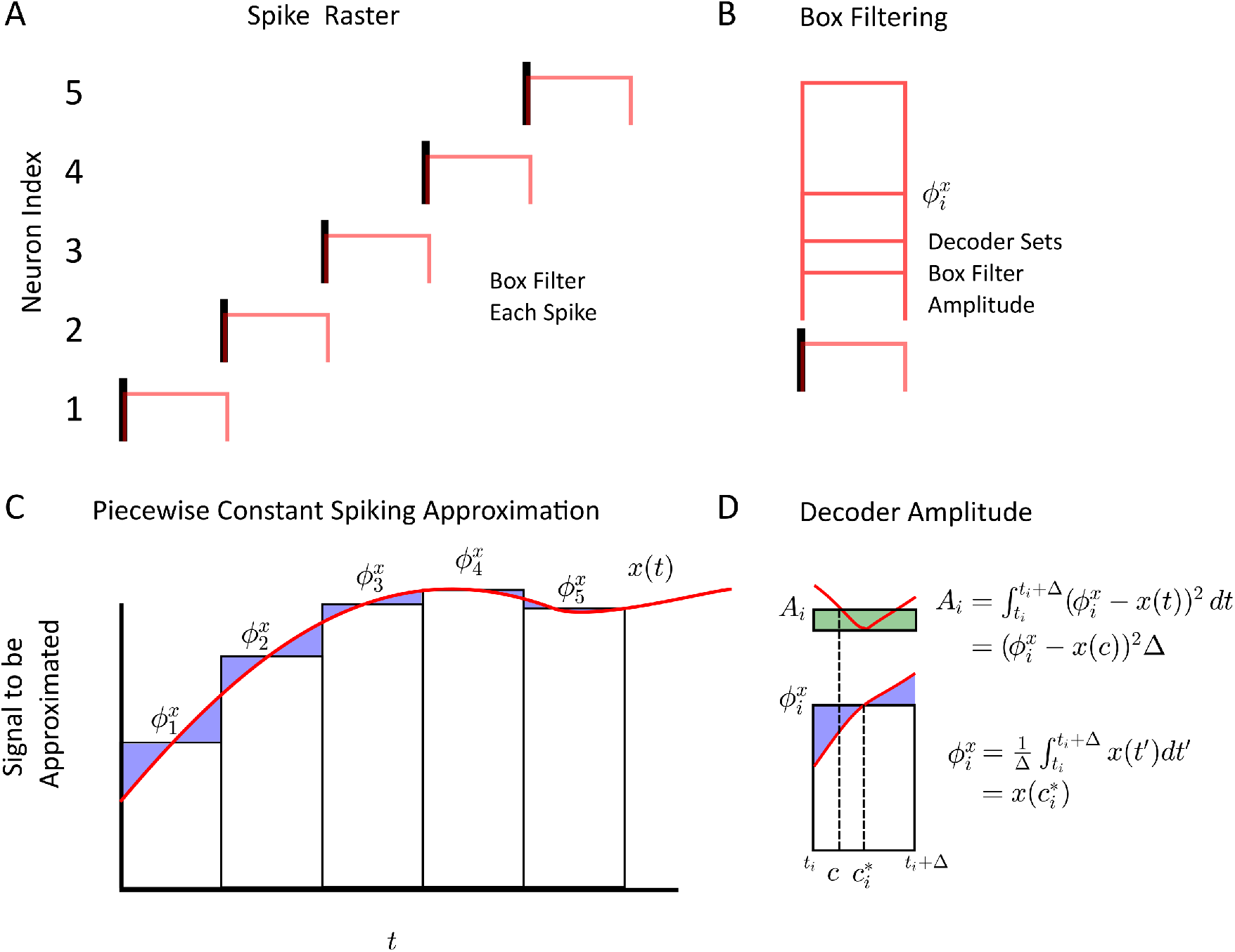
**(A)** The proof in the supplementary appendix for *O*(*N*^−1^) scaling relies on a chain of individual spikes which are box filtered to transform the inherently discrete spikes into a signal that covers a bin of time. **(B)** The optimal decoder for box-filtered, orthogonal spikes becomes the amplitude of each individual bin value. **(C)** The bins approximate continuous functions by setting the decoder value to the mean of the function value over the discrete-time bin a single spike covers. **(D)** Analytical determination of the decoder and application of the mean-value theorem for integrals.

**Supplementary Figure 2:**
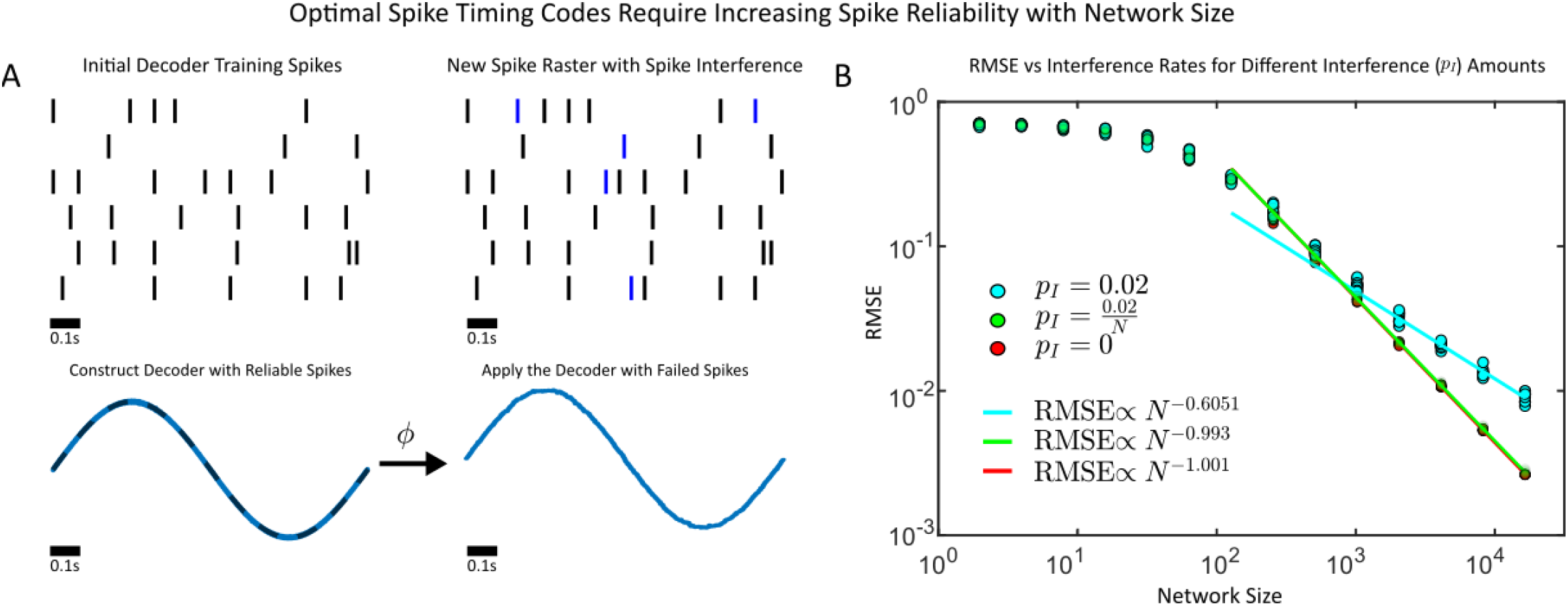
**(A)** The impacts of spike interference on decoding accuracy. (Top) In spike interference, spikes are randomly activated at segments of time where they were not expected by the trained decoder. (Bottom) The initial spike train is used to construct a decoder. Spikes are then randomly added with the same decoder applied, and the resulting error is measured for varying network sizes. **(B)** The root mean squared error (RMSE) for the decoded signal (sinusoidal oscillator) for networks with fixed spike interference (*p_F_* = 0.02, blue), no spike interference (*p_F_* = 0.00, red) and increasingly reliable spikes 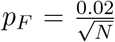 for larger networks, where *p_F_* is the probability that a spike is randomly added (see Materials and Methods).

